# Amygdala TDP-43 pathology is associated with behavioural dysfunction and ferritin accumulation in amyotrophic lateral sclerosis

**DOI:** 10.1101/2024.06.01.596819

**Authors:** Olivia M. Rifai, Fergal M. Waldron, Judi O’Shaughnessy, Fiona L. Read, Martina Gilodi, Annalisa Pastore, Neil Shneider, Gian Gaetano Tartaglia, Elsa Zacco, Holly Spence, Jenna M. Gregory

**Affiliations:** Centre for Discovery Brain Sciences, University of Edinburgh, UK; Department of Neurology, Center for Motor Neuron Biology and Disease, Columbia University, New York, USA; Institute of Medical Sciences, University of Aberdeen, UK; Department of Chemistry, University of Edinburgh, UK; RNA System Biology Lab, Center for Human Technology, Istituto Italiano di Tecnologia, Genoa, Italy; The Maurice Wohl Institute, King’s College London, London, UK

**Keywords:** ECAS, TDP-43, cognition, behaviour, ALS, Neuropathology

## Abstract

**Background:** Cognitive and behavioural symptoms associated with amyotrophic lateral sclerosis and frontotemporal spectrum disorders (ALSFTSD) are thought to be driven, at least in part, by the pathological accumulation of TDP-43.

**Methods:** Here we examine *post-mortem* tissue from six brain regions associated with cognitive and behavioural symptoms in a cohort of 30 people with sporadic ALS (sALS), a proportion of which underwent standardized neuropsychological behavioural assessment as part of the Edinburgh Cognitive ALS Screen (ECAS).

**Results:** Overall, the behavioural screen performed as part of the ECAS predicted accumulation of pathological phosphorylated TDP-43 (pTDP-43) with 100% specificity and 86% sensitivity in behaviour-associated brain regions. Notably, of these regions, pathology in the amygdala was the most predictive correlate of behavioural dysfunction in sALS. In the amygdala of sALS patients, we show variation in morphology, cell type predominance, and severity of pTDP-43 pathology. Further, we demonstrate that the presence and severity of intra-neuronal pTDP-43 pathology, but not astroglial pathology, or phosphorylated Tau pathology, is associated with behavioural dysfunction. Cases were also evaluated using a TDP-43 aptamer (TDP-43^APT^), which revealed that pathology was not only associated with behavioural symptoms, but also with ferritin levels, a measure of brain iron.

**Conclusions:** Intra-neuronal pTDP-43 and cytoplasmic TDP-43^APT^ pathology in the amygdala is associated with behavioural symptoms in sALS. TDP-43^APT^ staining intensity is also associated with increased ferritin, regardless of behavioural phenotype, suggesting that ferritin increases may occur upstream of clinical manifestation, in line with early TDP-43^APT^ pathology, representing a potential region-specific imaging biomarker of early disease in ALS.

**Key Messages:** *What is already known on this topic:* The amygdala is a key brain region in regulating behavior and emotional cognition and has been shown recently, through imaging studies, to be affected in ALS and FTD patients.

*What this study adds:* Here we examine the underlying pathology driving the association between the amygdala and behavioural symptoms in sporadic ALS demonstrating that region specific TDP-43 pathology and brain iron accumulation could represent potential early biomarkers of dysfunction.

*How this study might affect research, practice, or policy:* The correlation between early TDP-43 pathology (detected by RNA aptamer) and increased ferritin (brain iron accumulation) occurring upstream of clinical manifestation represents a potential, region-specific (amygdala), early imaging biomarker in ALS. This means that people at risk could be identified early and stratified for clinical trials prior to substantial neuronal cell loss and symptom onset.

## Introduction

Amyotrophic lateral sclerosis and frontotemporal spectrum disorders (ALSFTSD) is an umbrella term representing a genetically, pathologically, and clinically heterogeneous disease including people experiencing motor, cognitive and behavioural symptoms. As clinical trials are becoming more accessible to ALS patients, it is important to develop the ability to stratify patients for targeted therapies tailored to their predominating genetic, clinical, or pathological profile. Cognitive and behavioural dysfunction is a well-recognized consequence in up to 50% of patients with amyotrophic lateral sclerosis (ALS), with as many as 15% of ALS patients exhibiting frank frontotemporal dementia (FTD) (1). For this study, we were interested in interrogating pathological correlates of behavioural dysfunction in a cohort of sporadic ALS (sALS) patients. ALS with behavioural impairment (ALSbi) is defined by the presence of apathy or two other behaviour features, as per the consensus criteria (2). Symptoms of domain dysfunction included: (i) presence of apathy, (ii) behavioural disinhibition, (iii) loss of sympathy or empathy, (iv) perseverative and stereotyped behaviour or (v) hyperorality and change in eating habits. We have shown previously that pathological accumulation of phosphorylated TDP-43 (pTDP-43) in extra-motor brain regions is a specific, but not sensitive, pathological correlate of cognitive dysfunction that can be detected during life using the Edinburgh Cognitive ALS Screen (ECAS). The ECAS is a multi-domain, brief and extensively validated cognitive assessment tool that includes assessment of cognitive functions typically affected in ALS (i.e., executive function, social cognition, language and fluency) (3,4). The ECAS provides high sensitivity and specificity in the detection of even mild cognitive dysfunction in ALS against gold standard neuropsychological assessment using published cut-off scores for abnormality (2). The ECAS has been validated, as a sensitive screening tool for the detection of cognitive and behavioural dysfunction in many diverse populations of ALS patients and against other neuropsychological screening tools and batteries (5–11). However, this previous work has focused on pathological accumulation of pTDP-43 in cortical brain regions associated with motor and cognitive function. In our previous work, we did not have sufficient cases with behavioural dysfunction to draw firm conclusions about the relationship between behavioural dysfunction and pTDP-43 accumulation and have therefore since extended the cohort to separately evaluate this. Indeed, a sensitive pathological correlate of behavioural dysfunction is yet to be determined despite dysfunction of behaviour and emotional cognition being a much earlier and more sensitive clinical indicator of cortical dysfunction in ALS-Frontotemporal spectrum disorders (ALS-FTSD) (12).

Studies definitively assessing the clinico-pathological correlation of cognitive and behavioural dysfunction in ALS patients are often difficult due to differences in the neuropsychological tests performed during life, with many patients having undergone no neuropsychological testing at all (13, 14). Clinical imaging studies of patients with linked clinical data have identified multiple brain regions affected in ALS patients with behavioural dysfunction (15). Imaging of individuals with and without behavioural involvement, defined based on published criteria for ALSbi, has shown that patients with impaired behaviour had thinner bilateral precentral gyri, frontal, parietal, and temporal regions, and reduced volumes of the amygdala bilaterally, and enlarged inferior lateral and third ventricles. Therefore, in this study, we examined *post-mortem* tissue from six brain regions overlapping with the regions that showed imaging phenotypes in these ALSbi patients (15). The regions assessed included: (i) amygdala, (ii) orbitofrontal cortex (BA11/12), (iii) ventral anterior cingulate (BA24), (iv) medial prefrontal cortex (BA6), (v) prefrontal cortex (BA9) and (vi) the dorsolateral prefrontal cortex (BA46). Within this cohort of 30 sALS patients, a proportion underwent the same neuropsychological behavioural assessment as part of their ECAS evaluation. The aim of this study was to assess whether pTDP-43 pathology within these brain regions could explain associated imaging phenotypes and clinical manifestations of cognitive or behavioural symptoms as measured by the ECAS in our cohort. It is our hope that with a greater understanding of the clinico-pathological correlations underpinning extra-motor disease involvement, stratification of patients for trial inclusion and monitoring treatment response will become more effective and enable a greater breadth of treatment options to be available to patients in the future.

## Methods

### Cohort selection, clinical data and ethics

We selected all patients within the Edinburgh Brain Bank who had also undergone whole genome sequencing and/or gene panel assessment to establish that they had no underlying genetic mutation known to cause ALS (i.e., all sALS cases). All clinical data including the ECAS were collected as part of Scottish Motor Neuron Disease Register (SMNDR) and Care Audit Research and Evaluation for Motor Neuron Disease (CARE-MND) platform (ethics approval from Scotland A Research Ethics Committee 10/MRE00/78 and 15/SS/0216) and all patients consented to the use of their data during life. All *post-mortem* tissue was collected via the Edinburgh Brain Bank (ethics approval from East of Scotland Research Ethics Service, 16/ES/0084) in line with the Human Tissue (Scotland) Act. Use of human tissue for *post-mortem* studies has been reviewed and approved by the Edinburgh Brain Bank ethics committee and the Academic and Clinical Central Office for Research and Development (ACCORD) medical research ethics committee (AMREC).

### Neuropathological assessment

Brain tissue was taken at post-mortem from standardized Brodmann areas (BA) and fixed in 10% formalin for a minimum of 48 hours. Tissue was dehydrated in an ascending alcohol series (70-100%) followed by three successive four-hour washes in xylene. Three successive five-hour paraffin wax embedding stages were performed followed by cooling and sectioning of the formalin-fixed paraffin embedded (FFPE) tissue on a Leica microtome in 4 μm sections on to a superfrost microscope slide. Sections were dried overnight at 40°C and immunostaining was performed using the Novolink Polymer detection system with (i) the Proteintech anti-phospho(409–410)-TDP-43 antibody at a 1 in 4000 dilution; (ii) DAKO anti-GFAP antibody (Z0334) at a 1 in 800 dilution (iii) ThermoFisher anti-Tau antibody (cat no: MN1020) at a 1 in 2000 dilution (iv) Abcam Recombinant anti-Ferritin antibody (ab287968) at 1 in 100 dilution. DAB chromogen was used and counterstaining was performed with haematoxylin, according to standard operating procedures. Pretreatment for pTDP-43 was 30 minutes in a pressure cooker in pH6 buffered citric acid, pretreatment for Ferritin was 10 minutes in a pressure cooker in Tris/EDTA buffer and neither the pTau nor the GFAP antibodies required pretreatment. TDP-43^APT^ staining was performed as previously published (23).

pTDP-43, TDP-43^Apt^, GFAP and pTau pathology was graded manually by a trained histopathologist (JMG) using a semi-quantitative scoring system from 0-3: 0 = No pathology; 1 = Mild (1 to 5 affected cells in at least one 20x field of view per section); 2 = Moderate (5-15 affected cells in at least one 20x field of view per section); 3 = Severe (>15 cells affected in at least one 20x field of view per section). Cases defined as having no pathology had a score of 0, while those with pathology had a score from 1 to 3. This scoring was applied to neuronal and glial cell populations (based on established neuropathological knowledge of cell morphology using haematoxylin counterstain). The assessor was blinded to all demographic and clinical information. Digital burden scoring was performed using the freely available QuPath software implementing superpixel analysis using the following code: setImageType(’BRIGHTFIELD_H_DAB’); setColorDeconvolutionStains(’{“Name” : “H-DAB default“, “Stain 1” : “Hematoxylin“, “Values 1” : “0.65111 0.70119 0.29049 “, “Stain 2” : “DAB“, “Values 2” : “0.26917 0.56824 0.77759 “, “Background” : ” 255 255 255 “}’); setPixelSizeMicrons(0.625,0.625); createSelectAllObject(true); selectAnnotations (); runPlugin(’qupath.imagej.superpixels.DoGSuperpixelsPlugin’, ’{“downsampleFactor“: 1.0, “sigmaMicrons“: 3, “minThreshold“: 10.0, “maxThreshold“: 230.0, “noiseThreshold“: 0.0}’); selectDetections(); runPlugin(’qupath.lib.algorithms.IntensityFeaturesPlugin’, ’{“downsample“: 1.0, “region“: “ROI“, “tileSizePixels“: 200.0, “colorOD“: false, “colorStain1“: false, “colorStain2“: true, “colorStain3“: false, “colorRed“: false, “colorGreen“: false, “colorBlue“: false, “colorHue“: false, “colorSaturation“: false, “colorBrightness“: false, “doMean“: true, “doStdDev“: false, “doMinMax“: false, “doMedian“: false, “doHaralick“: false, “haralickDistance“: 1, “haralickBins“: 32}’); setDetectionIntensityClassifications(“ROI: 2.00 µm per pixel: DAB: Mean“, 0.2689, 0.3007, 1) as done previously (20). Number of 1+ detections and more intense 2+ detections were quantified and the weighted score (1x number of 1+ detections + 2x number of 2+ detections) was compared to manual scoring of estimated pTau neuropil burden to ensure adequate quality control. Graphs were plotted using R. Fisher’s exact test and Mann Whitney U tests were used as appropriate to the data distribution and stated in the figure legends.

## Results

### Behavioural dysfunction detected by the ECAS is associated with pathological TDP-43 accumulation

Thirty sALS patients were assessed for the severity and cell-type specificity of TDP-43 aggregates using the phospho-TDP-43 antibody, which stains only pathological TDP-43 aggregates. Pathology was graded as absent (0), mild (1), moderate (2) or severe (3) for both neuronal inclusions and glial inclusions (Figure 1A; Table 1). Out of the 30 patients, 12 had undergone behavioural evaluation during life, with 4 meeting criteria for ALSbi. In total, 6 patients had behavioural domain dysfunction and 6 had no evidence of any behavioural involvement. The frequency of different behavioural symptoms is detailed in Figure 1B, with perseveration and apathy being the most common symptoms in our cohort. The overall assessment showed that all cases with a detectable behavioural or cognitive domain affected had pathological pTDP-43 present in the selected brain regions. Overall, the presence of pTDP-43 in these six brain regions, when evaluated together, was associated with both cognitive dysfunction (i.e., ECAS score below published cut offs for at least one domain out of fluency, executive and language; *p* = 0.0137) and behavioural dysfunction (*p* = 0.0152; Figure 1C).

**Figure 1.**
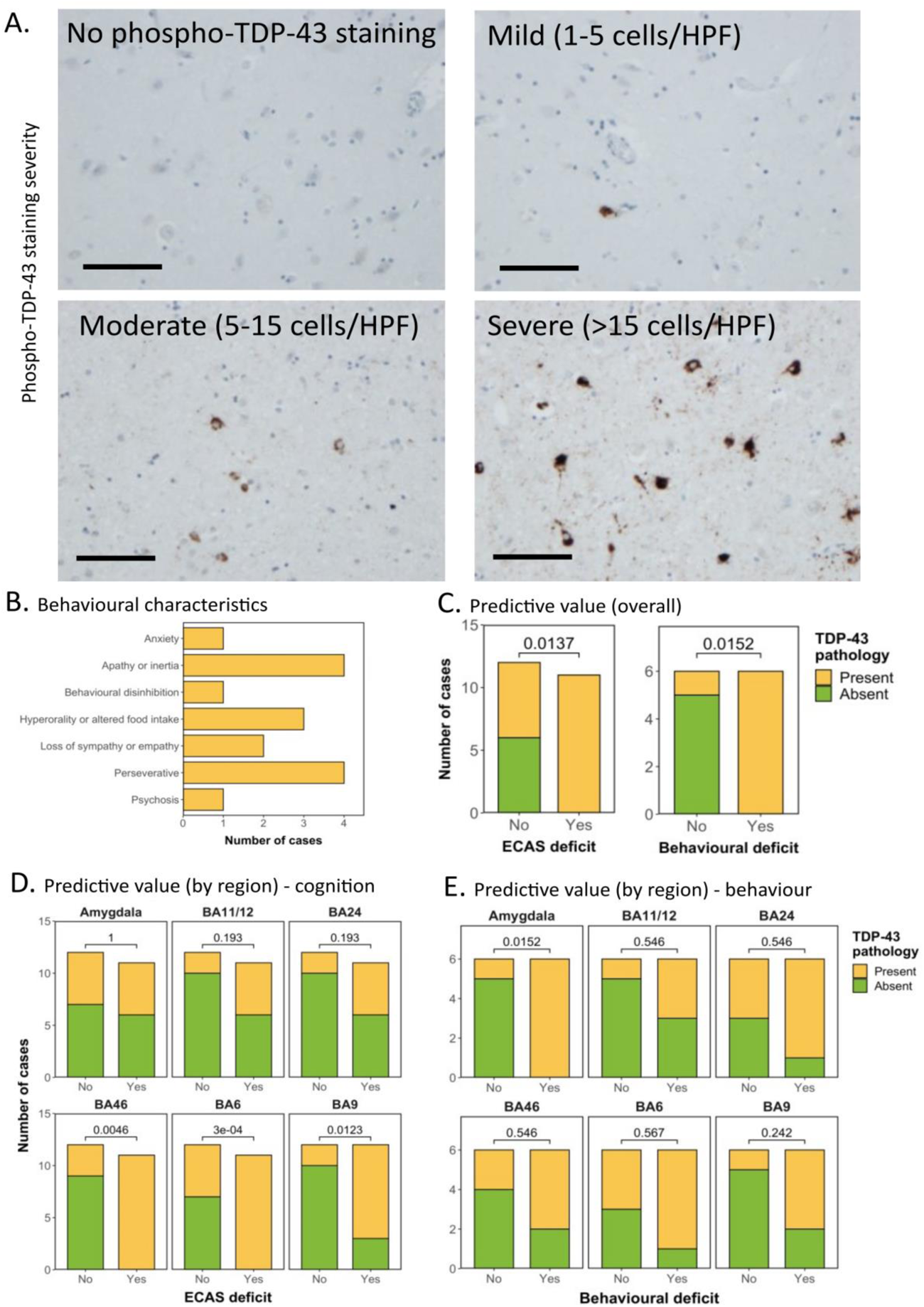
Behavioural dysfunction detected by the ECAS is associated with pathological TDP-43 accumulation. A. Representative photomicrographs taken at 20 x magnification demonstrating the varying severity of phospho-TDP-43 pathology in the amygdala in sALS cases. Pathology ranges from completely absent to severe (>15 cells per high power field (HPF; scale bar = 100 µm). **B.** Frequency distribution demonstrating the behavioural domains affected in the 6 patients with behavioural dysfunction detected by ECAS, demonstrating that perseveration and apathy are the most affected domains. **C.** Graph detailing the association between the absence/presence of pTDP-43 in all six brain regions overall, and the absence/presence of cognitive dysfunction (left) or behavioural dysfunction (right). Demonstrating that the presence of pTDP-43, using all six regions combined, has a significant predictive value for the detection of both cognitive and behavioural dysfunction. **D&E.** Graph detailing the association between the absence/presence of pTDP-43 in all six brain regions individually, and the absence/presence of cognitive dysfunction (**D**) or behavioural dysfunction (**E**). Demonstrating that the presence of pTDP-43, in BA46, BA6 and BA9, has a significant predictive value for cognitive dysfunction and the presence of pTDP-43, in the amygdala has a significant predictive value for behavioural dysfunction.

**Table 1.**
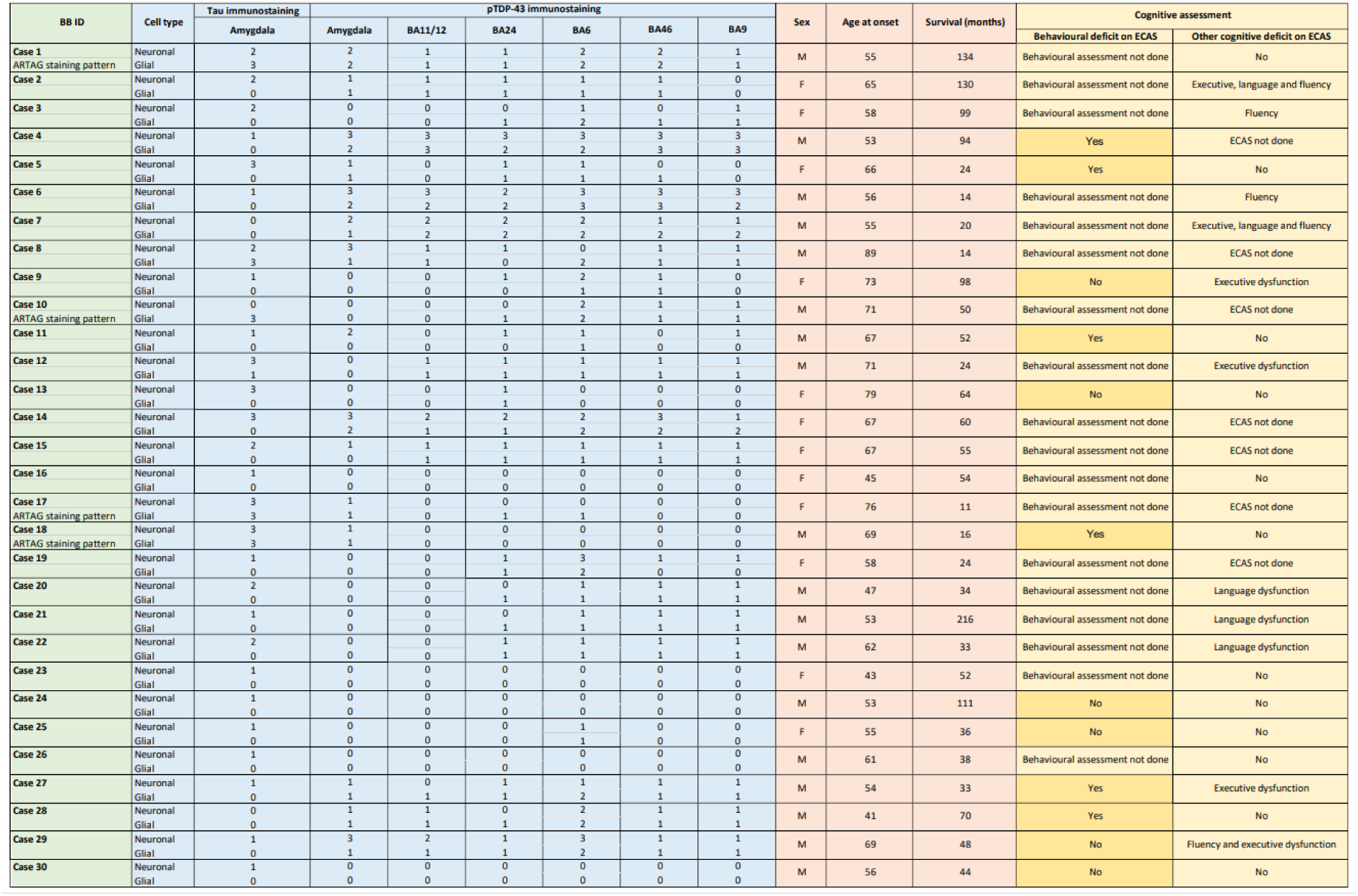
Cohort demographics. Table detailing cohort demographics including Tau and TDP-43 severity scoring, clinical and cognitive data.

### Amygdala pTDP-43 pathology is the most sensitive correlate of behavioural dysfunction

We next sought to establish whether there were specific brain regions driving these effects or whether the predictive value was in assessing all regions together. The brain regions with the strongest association with cognitive dysfunction were BA46, BA6 and BA9, frontal brain regions typically associated with executive function (Figure 1D). The amygdala was the only brain region assessed that was associated with behavioural dysfunction (Figure 1E). No specific predictive value was found for all remaining regions (Figure 1E).

### Variation in cell-type predominance and severity of phospho-TDP-43 immunostaining in the amygdala of sALS patients

Given that the presence of pTDP-43 pathology in the amygdala was the most sensitive pathological correlate of behavioural dysfunction in our sALS cohort, we next sought to further characterize pTDP-43 aggregates and pathological features in this region. Of the 30 sALS cases profiled, 15 demonstrated TDP-43 pathology in the amygdala, 8 of which exhibited a predominantly neuronal pattern of staining (Figure 2A; left) and 7 of which exhibited a mixed glial and neuronal staining pattern (Figure 2A; right). Interestingly, we previously reported a pure, glial-only staining pattern in 22.2 % of ALS cases (4) in other cortical brain regions, which was not observed in the amygdala in this sALS cohort. Furthermore, no cases exhibited severe glial TDP-43 pathology and only a small number (n = 4) exhibited moderate glial pathology. Most cases had either no glial (n = 17) pathology or mild glial pathology (n = 9). This is in contrast with neuronal TDP-43 pathology, for which mild (n = 7), moderate (n = 3) and severe (n = 5) pathologies existed more frequently, with 15 cases showing no neuronal TDP-43 pathology (Table 1). Furthermore, astroglial aggregates observed in the amygdala of all cases were morphologically distinct to those seen in laminated cortical regions such as the motor cortex (Figure 2B), with cortical glia exhibiting single, compact amorphous aggregates in a perinuclear distribution and amygdala glia exhibiting multiple aggregates, ranging in size and occupying perinuclear regions as well as proximal processes.

**Figure 2.**
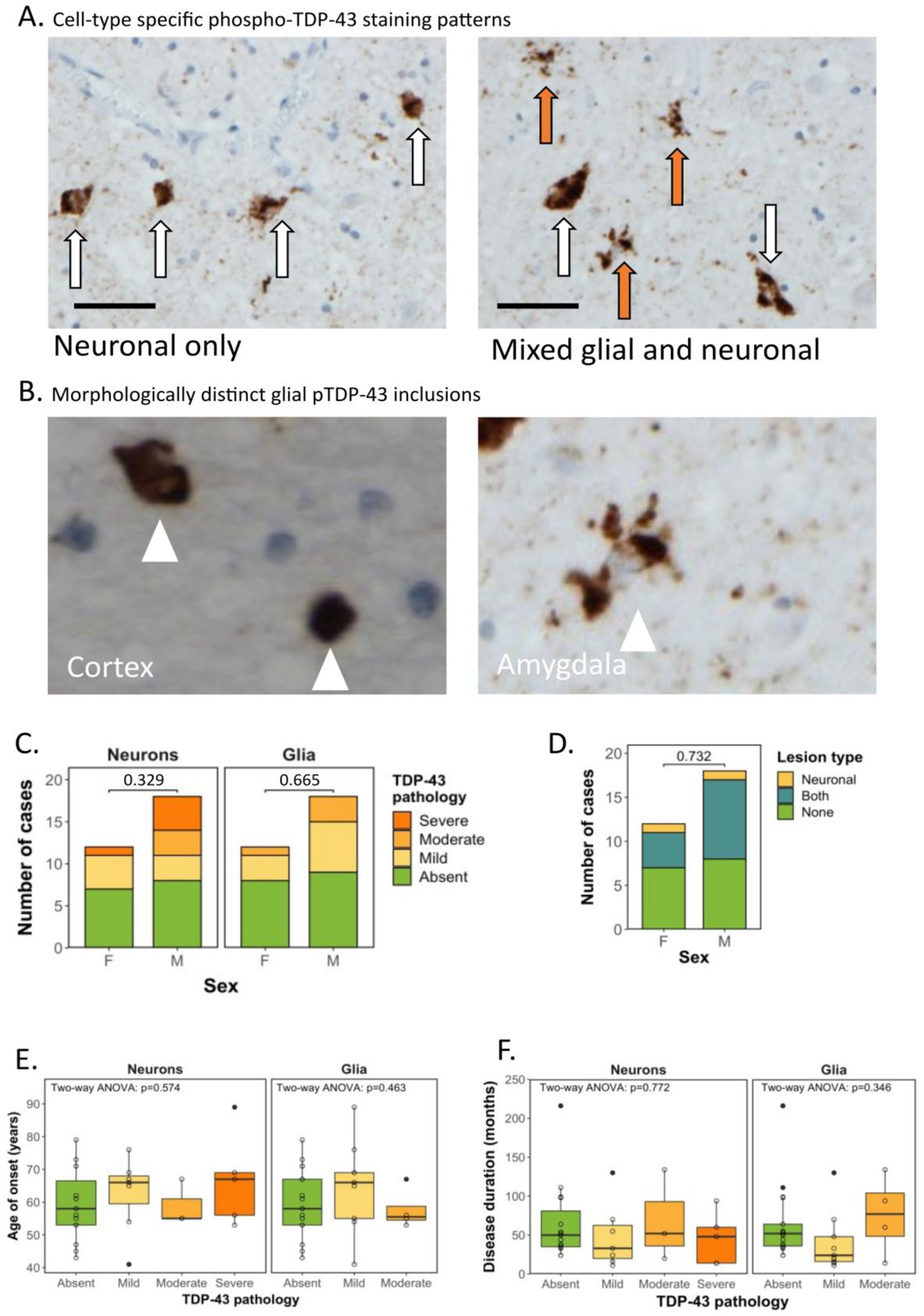
Variation in cell-type predominance and severity of pTDP-43 immunostaining in the amygdala of sALS patients. **A.** Representative photomicrographs taken at 40x magnification demonstrating cell-type specific predominance of phospho-TDP-43 pathology in the amygdala in sALS cases. Brown (DAB) staining is phospho-TDP-43, blue counterstain is haematoxylin. Left: TDP-43 staining in a neuronal-only pattern (scale bar = 50 µm; white arrows indicate neurons containing pathological TDP-43 aggregates). Right: TDP-43 staining in a mixed neuronal and glial pattern (scale bar = x µm; white arrows indicate neurons and red arrows indicate glial cells containing pathological TDP-43 aggregates). **B.** Representative high power photomicrographs demonstrating glial aggregates (white arrowheads) with a distinct morphology in cortical regions (left) exhibiting single, compact amorphous aggregates in a perinuclear distribution compared with glia in the amygdala (right) exhibiting multiple aggregates, ranging in size occupying perinuclear regions as well as proximal processes. **C.** Graph comparing the presence and severity of neuronal TDP-43 pathology (left), glial TDP-43 pathology (centre) and **D**. type of lesion (neuronal, glial or mixed TDP-43 pathology) to sex of patient demonstrating no association tested by Fisher’s exact test (p = 0.329, p = 0.665 and 0.732 respectively). **E.** Graph comparing neuronal TDP-43 severity (left) and glial TDP-43 severity (right) to age of disease onset demonstrating no association (Two-way ANOVA). **F.** Graph comparing neuronal TDP-43 severity (left) and glial TDP-43 severity (right) to disease duration (months from disease onset to death) demonstrating no association (Two-way ANOVA).

### No association between clinical characteristics and pTDP-43 presence or severity in the amygdala

Assessing the relationship between pTDP-43 pathology and clinical characteristics, Fisher’s exact test revealed no statistically significant relationship between sex and the severity of neuronal (*p* = 0.329) or glial TDP-43 pathology (*p* = 0.665; Figure 2C), nor between sex and the type of lesion present (i.e. neuronal, mixed (both) or no pathology; *p* = 0.732; Figure 2D). As pTDP-43 accumulation has been associated with alterations in amygdala shape and volume in healthy ageing (16), we next assessed whether the differential burden in amygdala pTDP-43 pathology is related to age of disease onset. Age of onset had no significant relationship with neuronal (*F*(3,257) = 0.677, *p* = 0.574) or glial (*F*(2,197) = 0.793, *p* = 0.463) pTDP-43 severity (Figure 2E). We next hypothesized that accumulation of TDP-43 pathology could be a function of disease duration, with longer disease duration allowing for prolonged accumulation of pathology. However, disease duration had no significant relationship with neuronal (*F*3,2461) = 0.375, *p* = 0.772) or glial (*F*(2,4489) = 1.103, *p* = 0.346) pTDP-43 severity (Figure 2F).

### Presence and severity of intra-neuronal pTDP-43 pathology, but not astroglial or pTau pathology, is associated with a clinically detectable behavioural manifestation

Out of the 30 sALS patients that we evaluated, 12 had undergone behavioural evaluation during life, with 6 meeting criteria for ALSbi and 6 with no evidence of any behavioural involvement. Of the 12 clinically profiled sALS patients in our cohort, if there was evidence of behavioural dysfunction, these individuals always exhibited pTDP-43 pathology in the amygdala (n = 6), similarly to our previous findings in other non-motor brain regions (4). We also observed that if TDP-43 pathology was not present, there was no evidence of behavioural dysfunction (n = 5), except for a single case, which had no evidence of behavioural dysfunction despite significant pTDP-43 burden (Table 1). However, this patient did have a recorded entry of psychosis in their medical history, which was well controlled with treatment and not detected in the psychosis section of the ECAS behavioural screen. This is in keeping with our previous reports of mismatch cases in ALS, and mismatch cases that exist for other neurodegenerative diseases, in which TDP-43 burden is not related to clinical manifestations of dysfunction in those brain regions (4).

We next sought to establish whether cell type specific TDP-43 pathology in the amygdala was related to behavioural symptom burden. Fisher’s exact test revealed a statistically significant relationship between the severity of neuronal TDP-43 pathology and detectable behavioural dysfunction, with more severe neuronal TDP-43 pathology being associated with the presence of behavioural dysfunction (Figure 3A; *p* = 0.006). This was not the case for the severity of glial TDP-43 pathology, which showed no statistically significant relationship with the presence of behavioural dysfunction (Figure 3A; *p* = 0.080). Given that previous reports examining the relationship between TDP-43 pathology and cognitive correlates have only examined absence versus presence of pathology rather than severity, we also examined absence versus presence. Fisher’s exact test revealed that there was a statistically significant association between presence of behavioural dysfunction and neuronal pathology (*p* = 0.015) but not for glial pathology (*p* = 0.08) (Figure 3A).

**Figure 3.**
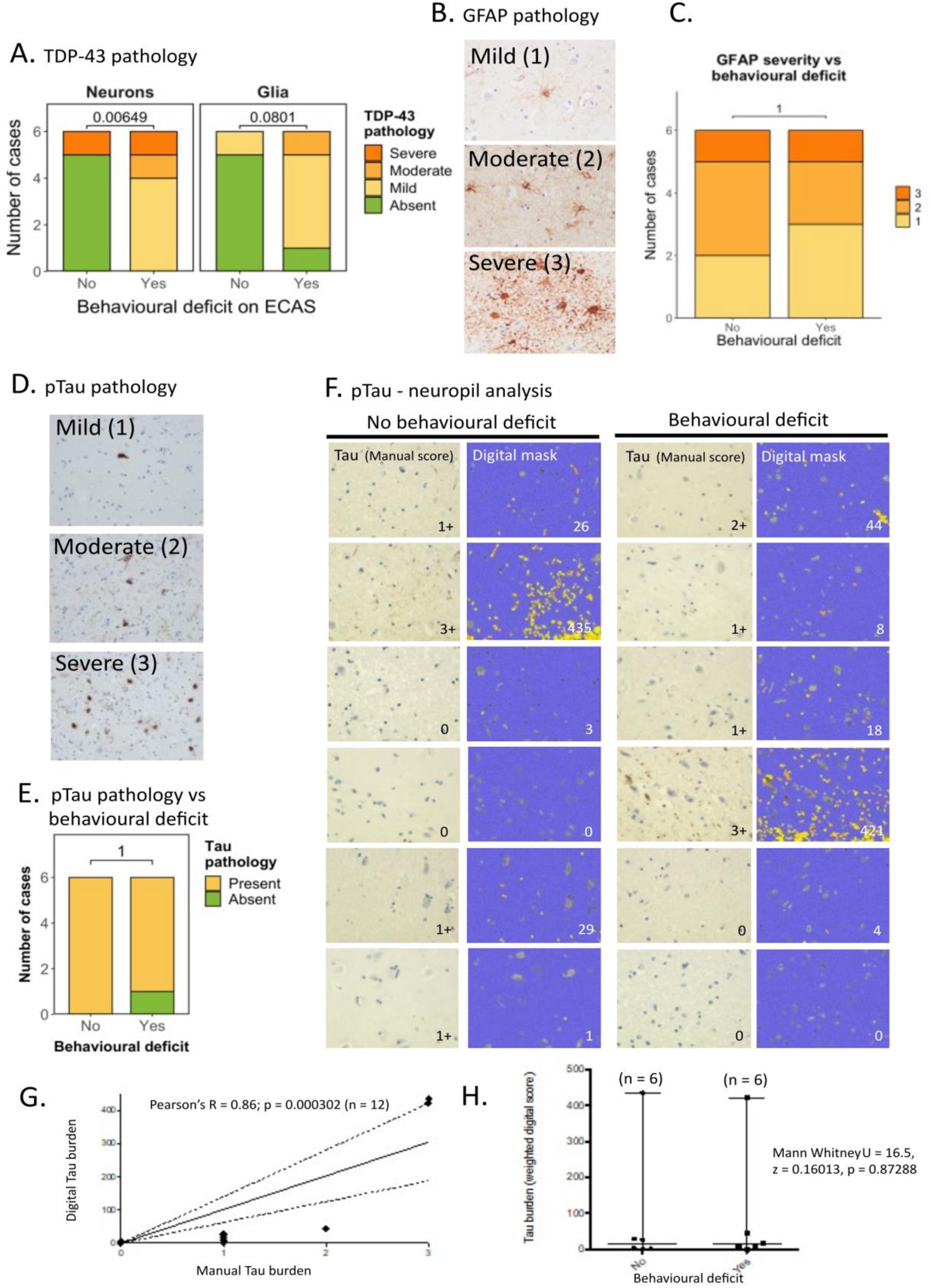
Presence and severity of intra-neuronal pTDP-43 pathology, but not astroglial or Tau pathology, is associated with a clinically detectable behavioural manifestation. **A.** (Left) Graph comparing the presence and severity of neuronal pTDP-43 pathology to the presence of clinically detectable behavioural dysfunction. (Right) Graph comparing the presence and severity of glial TDP-43 pathology to the presence of clinically detectable behavioural dysfunction. These data demonstrate an association with the severity of neuronal TDP-43 (p = 0.006) but not the severity of glial TDP-43 (p = 0.080) tested by Fisher’s exact test. Statistical examination of the absence or presence of TDP-43 pathology vs the absence or presence of behavioural dysfunction also results in an association with the presence of neuronal TDP-43 (p = 0.015) but not the presence of glial TDP-43 (p = 0.08) tested by Fisher’s exact test. **B.** Representative photomicrographs taken at 40 x magnification demonstrating the varying severity of glial reactivity using GFAP immunohistochemical staining in the amygdala in sALS cases. Pathology ranges from mild to severe (scale bar = 50 µm). **C.** Graph comparing the severity of glial activation to the presence of clinically detectable behavioural dysfunction. These data demonstrate no association between the severity of glial reactivity and behavioural dysfunction tested by Fisher’s exact test (p=1.00). **D.** Representative photomicrographs taken at 20 x magnification demonstrating the varying severity of Tau pathology in the amygdala in sALS cases. Pathology ranges from mild to severe (HPF= high power field; scale bar = 100 µm). **E.** Graph comparing the presence and severity of Tau pathology to the presence of clinically detectable behavioural dysfunction (above) and cognitive dysfunction (below), demonstrating no statistically significant association. **F.** Representative photomicrograph from each of the 12 cases (n = 6 with behavioural dysfunction and n = 6 without) of amygdala stained with Tau AT8 antibody (left image) with adjacent digital mask generated implementing a superpixel analysis of DAB intensity using QuPath digital pathology analysis software. Yellow mask indicates intensity 1 staining and orange mask indicates stronger intensity 2 staining. Grey indicates manually excluded cell bodies to ensure the analysis of only neuropil. Blue indicates negative background. **G.** Correlation between weighted digital quantification on y-axis, with manual analysis of grading of neuropil Tau burden (0=absent, 1=mild, 2=moderate, 3=severe) on x-axis as part of digital quality control showing a statistically significant correlation between the two measures (Pearsons R = 0.8629, p = 0.000302). **H.** Mann Whitney U test shows no association between digitally quantified Tau neuropil burden and behavioural dysfunction (U = 16.5, z = 0.16013, p = 0.87288). Each case is indicated by a single character and median and range are plotted on the graph (n = 6 in each group).

Studies from other amygdala-associated diseases, such as temporal lobe epilepsy and progressive supranuclear palsy, have reported astrogliosis as key factor influencing disease susceptibility (17,18). As we did not see a significant association between glial TDP-43 pathology and the presence of behavioural dysfunction, we hypothesized that glial reactivity, rather than glial TDP-43 accumulation, may be more closely associated with behavioural dysfunction in these cases. We performed immunohistochemical staining for GFAP, a marker of astrogliosis (Figure 3B), and graded pathology as mild (1), moderate (2) or severe (3). Fisher’s exact test revealed no significant association between glial reactivity and the presence of behavioural dysfunction (*p* = 1.00) (Figure 3C).

Given the prevalence of co-pathologies (i.e., concomitant Tau and pTDP-43 pathology) previously reported in the amygdala of patients with frontotemporal lobar degeneration (FTLD) (19), we also characterised phospho-Tau (pTau) burden in our sALS cohort using the AT8 monoclonal antibody (Figure 3). We first scored pTau accumulation using a semi-quantitative scoring method (i.e., manual pathological analysis) to quantify the cell-type specific burden of pTau (Figure 3D). Interestingly, 28 out of the 30 sALS cases exhibited pTau pathology, 14 with mild pathology, 5 with moderate pathology and 9 with severe pathology. In 4 of these cases, we also observed aging-related pTau astrogliopathy (ARTAG)-like pathological features such as thorn astrocytes and perivascular/subpial distribution of staining. Whilst these features were observed in older individuals within our cohort, there were no other clinical features that stood out as correlates of ARTAG-like pathology (Table 1). These features have been recorded in Table 1 with associated clinical data, in case this may be useful for future meta-analysis to understand the clinico-pathological relevance of ARTAG-like pathology in neurodegenerative diseases when larger datasets become available. Two cases had pure pTDP-43 pathology, 15 cases had pure pTau pathology and 13 cases had overlapping pTDP-43 and pTau pathology. There was no significant association between pTau pathology in the amygdala and behavioural or any other ECAS impairment (Figure 3E).

Noting that pTau can accumulate in a neuropil-predominant fashion in some of the cases in our cohort, and that our cell-type specific quantification may underestimate this burden, we next performed digital analysis to quantify neuropil staining in cases with and without behavioural dysfunction. We hypothesized that pTau accumulation within the neuropil may be more indicative of pTau accumulation in processes and synapses and thus more predictive of cognitive dysfunction (Figure 3F). We first performed a semi-quantitative manual burden scoring estimate to validate our digital superpixel analysis as part of our digital analysis quality control (Figure 3G). Correlation between the digital superpixel analysis and the manual scoring was significant (Pearson’s correlation: *R* = 0.8629; *p* = 0.000302). Comparative analysis of pTau burden between behaviourally affected and unaffected individuals was then performed using Mann Whitney U, showing no significant difference between groups (*U* = 16.5; *z* = 0.16013; *p* = 0.87288; Figure 3H).

### TDP^APT^ staining reveals early nuclear TDP-43 pathology in discordant cases

We further characterised TDP-43 pathology in our cohort using our recently developed TDP-43 RNA aptamer (TDP^APT^) (23, 24). We have shown previously that TDP^APT^ has improved sensitivity compared to the pTDP-43 antibody and is able to detect aggregation events that are either missed (i.e., by the pTDP-43 antibody) or obscured (i.e., by the C-terminal TDP-43 antibody) (23). Using TDP^APT^, we show that all 6 cases with behavioural impairment exhibit extensive neuronal TDP-43 pathology within the nucleus and cytoplasm (Figure 4). Further, we demonstrate that 3 out of the 6 cases with no evidence of behavioural dysfunction also exhibit TDP-43^APT^ pathology (Figure 4). In these cases, there is prominent nuclear pathology (i.e., nuclear rods, nuclear membrane pathology and nucleolar decoration). Importantly, we have previously reported that these features are observed in early stages of TDP-43 pathology that precede clinical manifestation (23). The remaining 3 cases (Cases 9, 13, and 25) show no evidence of TDP-43 pathology or behavioural dysfunction.

**Figure 4.**
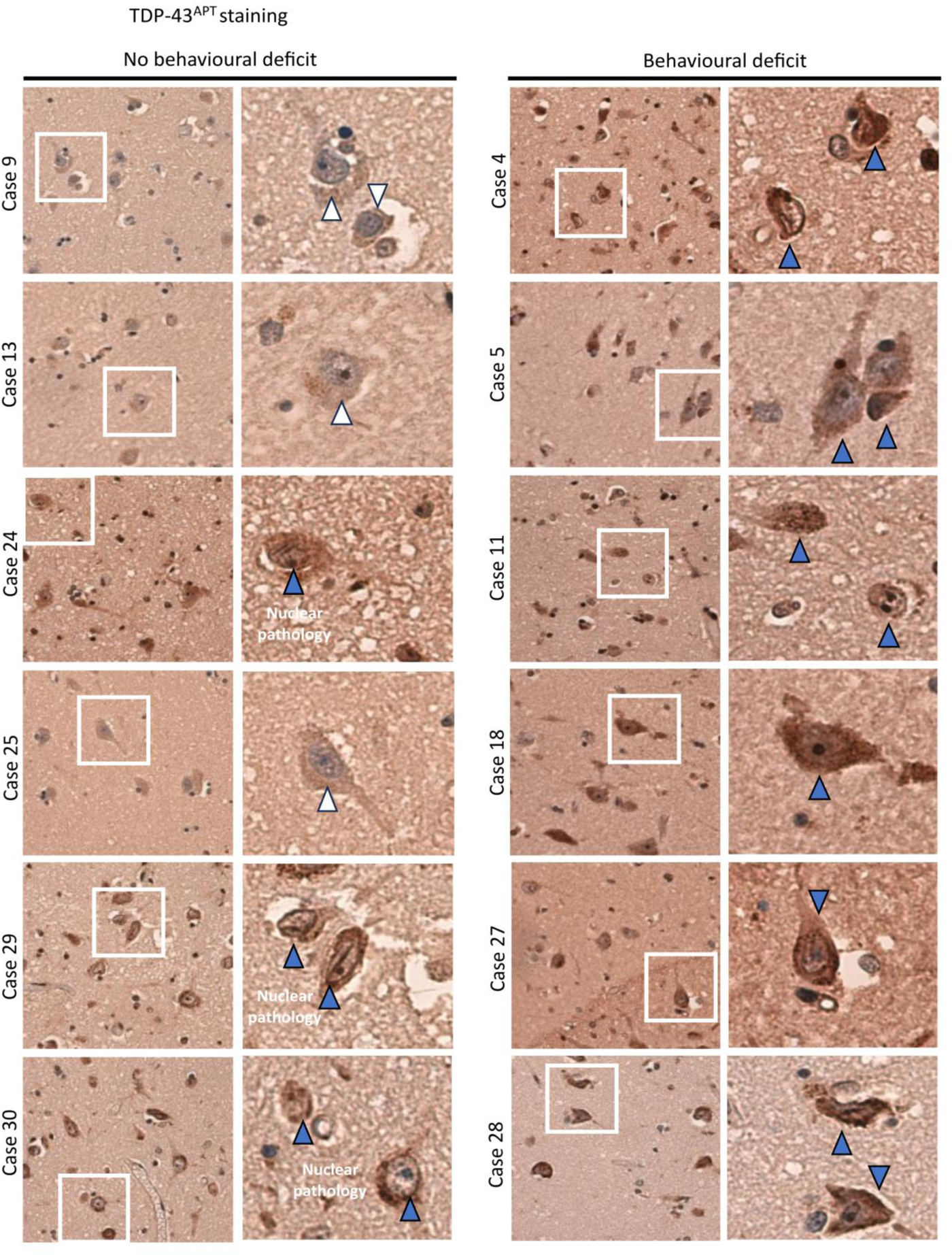
TDP^APT^ staining reveals early nuclear TDP-43 pathology in discordant cases. Representative photomicrographs taken at 20 x magnification demonstrating TDP-43^APT^ pathology in the amygdala in sALS cases. Cases with no evidence of a behavioural deficit are included on the left and those with a behavioural deficit on the right. Each panel comprises a low power image on the left and a digital zoom, of the area highlighted by the white box, on the right. White arrows indicate neurons with no evidence of TDP-43^APT^ pathology and blue arrows indicate neurons with TDP-43^APT^ pathology (including prominent nuclear pathology in non-symptomatic individuals and both nuclear and cytoplasmic pathology in symptomatic individuals).

### Ferritin levels correspond with TDP^APT^ pathology and clinical phenotype

To gain mechanistic insight into how and why TDP-43 aggregation occurs in the amygdala, and in turn how this may drive clinical phenotype in some but not all individuals, we next examined whether brain iron accumulation was associated with regional clinical phenotype. We hypothesized that brain iron accumulation, which has been reported in the amygdala of people with ALS, could precipitate protein peroxidation changes leading to protein aggregation. We thus examined the burden of iron accumulation in the form of ferritin in our cohort. Cases 9, 13, and 25, which showed no evidence of TDP-43 pathology in the amygdala, exhibited a cell-type specific ferritin staining pattern, predominantly restricted to the peri-nuclear region of glial cells (Figure 5A). In cases that had demonstrable TDP-43^APT^ pathology (either nuclear or established nuclear and cytoplasmic pathology) there was evidence of a shift in the subcellular localization of the ferritin staining, with prominent staining visible in the cell body and processes of glial cells and overall, a more intense staining pattern. We quantified DAB staining intensity digitally using superpixel analysis as described above. There was a significant correlation (Pearson’s correlation: *R* = 0.7; *p* = 0.0007) between ferritin and TDP-43^APT^ staining burden (Figure 5B). This correlation was specific to TDP-43 pathology, as we saw no significant correlation (Pearson’s correlation: *R* = 0.05; *p* = 0.5) between ferritin and pTau staining burden (Figure 5C). Mean ferritin intensity was also significantly increased in behaviourally impaired compared to unimpaired individuals (unpaired *t*-test, *p* = 0.022; Figure 5D).

**Figure 5.**
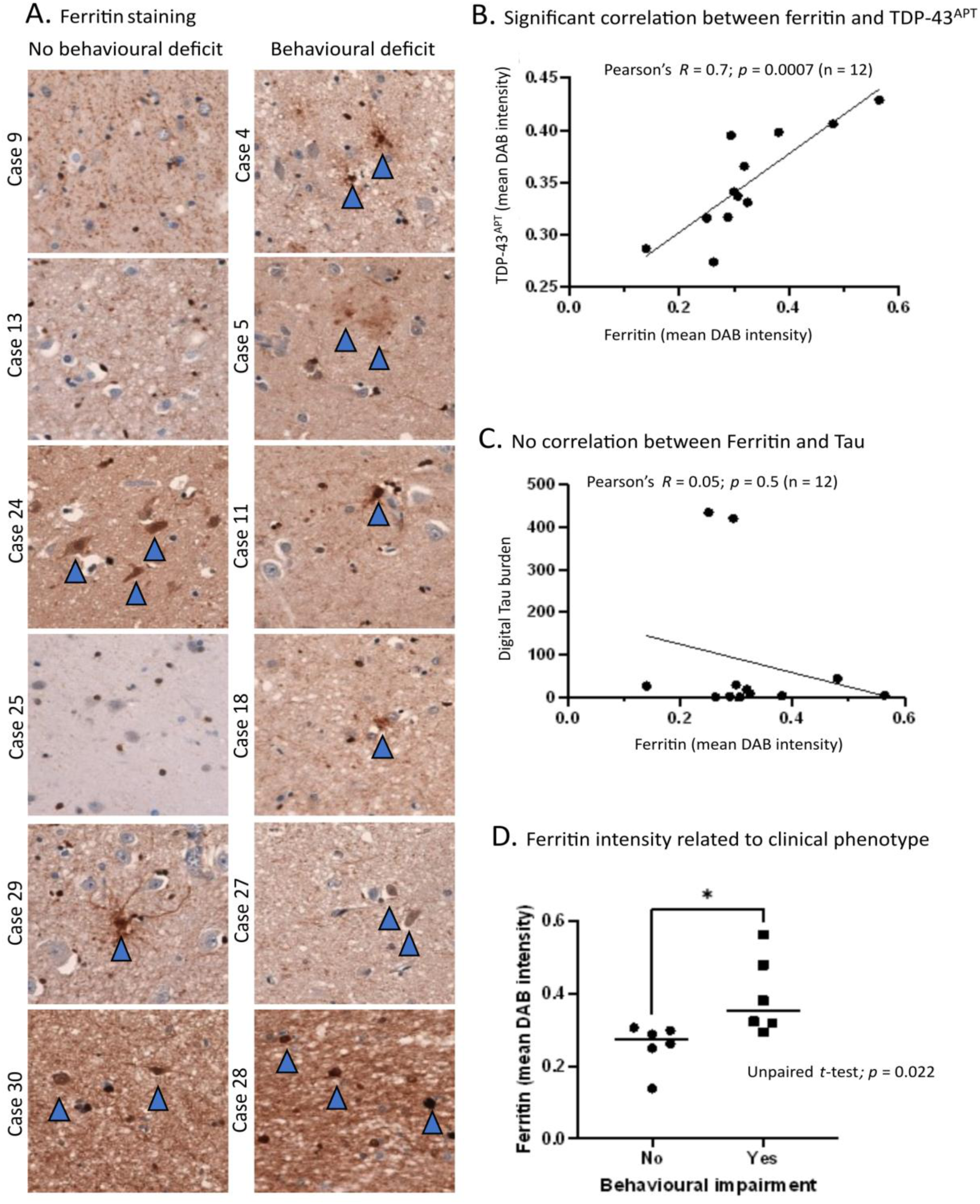
Ferritin levels correspond with TDPAPT pathology and clinical phenotype. **A**. Representative photomicrographs taken at 20 x magnification demonstrating TDP-43^APT^ pathology in the amygdala in sALS cases segregated by the presence (left panel) and absence (right panel) of behavioural deficits. Blue arrows indicate cells with pathological staining. **B**. Graph comparing digitally quantified mean DAB intensity for TDP-43^APT^ pathology on the y-axis and ferritin on the x-axis demonstrates a significant correlation (n=12). **C**. Graph comparing digitally quantified mean DAB intensity for Tau pathology on the y-axis and ferritin on the x-axis demonstrates no evidence of correlation (n=12). D. Graph demonstrating mean DAB intensity for each case stained with ferritin digitally acquired using superpixel analysis. Cases are grouped by behavioural deficit and an unpaired t-test demonstrates a statistically significant difference in ferritin staining between clinically distinct groups, with higher mean ferritin intensity noted in clinically symptomatic cases.

## Discussion

Here, we set out to understand the pathological correlates of behavioural dysfunction in ALSbi. We selected a cohort of 30 sALS patients and examined brain areas, identified from imaging studies of ALSbi patients, that are thought to be associated with behavioural dysfunction: the (i) amygdala, (ii) orbitofrontal cortex (BA11/12), (iii) ventral anterior cingulate (BA24), (iv) medial prefrontal cortex (BA6), (v) prefrontal cortex (BA9) and (vi) the dorsolateral prefrontal cortex (BA46). We demonstrated that the behavioural screen done as part of the ECAS was predictive of TDP-43 pathology in these regions with 100% specificity and 86% sensitivity. Regional analysis revealed that, out of the regions profiled, TDP-43 pathology in BA9, BA46 and BA6 had significant associations with cognitive dysfunction (as we have demonstrated previously; 4). Further, the amygdala was the only region profiled with a significant association between TDP-43 pathology and behavioural dysfunction.

The amygdala is involved early in many common forms of misfolded proteinopathy (21), and TDP-43 pathology has previously been reported in the amygdala of ALS patients (13, 14). However, a comprehensive analysis of TDP-43 pathology in the amygdala of a deeply clinically phenotyped cohort of ALS cases has, to date, not been carried out. We therefore examined *post-mortem* tissue from the amygdala in our cohort of 30 sALS patients, reporting variation in morphology, cell-type predominance, and severity of TDP-43 pathology in the amygdala. Previous analysis of TDP-43 pathology within the amygdala has demonstrated TDP-43 pathology in 38.5% of ALS-FTSD cases and an increased prevalence of pTDP-43 pathology in cases with dementia; however, many cases lacked adequate and/or comparable clinical information, limiting further conclusive analysis (14). As individuals in our cohort had been assessed using the same cognitive screening tool, we were able to examine associations between regional pTDP-43 pathology and ECAS subdomain scores for behaviour and emotional cognition. Employing this approach, we showed that presence and severity of intra-neuronal, but not glial, TDP-43 pathology in the amygdala was associated with behavioural dysfunction. Taken together, our data suggest that behavioural impairment detected using the ECAS is associated with pTDP-43 pathology in the amygdala and that neuronal TDP-43 pathology in the amygdala is a stronger correlate of behavioural dysfunction than glial pathology. By contrast, we and others have previously shown that the absence or presence of pTDP-43 pathology, but not cell-type specificity or severity of pTDP-43 pathology, is associated with cortical (i.e., motor and cognitive) dysfunction in ALS-FTSD (4, 22).

Our findings demonstrate both regional and cell-type specific predominance of intra-neuronal amygdala TDP-43 accumulation, the abundance of which is significantly associated with ferritin accumulation and clinically detectable behavioural dysfunction. Notably, mismatch cases were also identified, which, despite having intra-neuronal TDP-43 aggregate burden, did not have corresponding behavioural dysfunction. These observations were consistent with the frequency of unaffected mismatch individuals assessed in other cohorts (14). Furthermore, mismatch cases demonstrated early-stage, nuclear TDP-43 pathology (nuclear pathology only), detected by TDP-43^APT^, which we have demonstrated to be an early pathological feature that precedes clinical manifestation (23). Importantly, this is in line with reports that there is an increasing frequency of cognitive and behavioural symptoms with increasing disease stage in ALS (25).

Our observation of an increased intra-neuronal predominant pattern of pTDP-43 aggregation in ALSbi and lack of a significant association between behavioural impairment and glial pTDP-43 pathology (Figure 3A), glial reactivity (Figure 4B) or pTau pathology (Figure 4D) in the amygdala, suggests a selective, regional, cell-type specific vulnerability to TDP-43 associated neurodegeneration unlike in frontal cortex regions,. Further intra-regional pathophysiological variation is also suggested by the atrophy of specific amygdala nuclei on imaging (26). Indeed, other ECAS domains are highly predictive of brain region abnormalities on imaging (27, 28). We have also previously shown that the ECAS, performed during life, is highly specific (100% specificity) at identifying individuals with pTDP-43 pathology at *post-mortem*. The sensitivity of the ECAS is much lower, around 20-50% depending on the brain regions involved, owing to the incidence of discordant cases, which we have also identified in this analysis. As we have shown previously (23) and in this study (Figure 4) that discordant cases exhibit early TDP-43 pathological features that precede the clinical inflection point, this could make these features ideal for identifying people in early disease states.

We also report a significant positive correlation between TDP-43^APT^ and ferritin immunoreactivity, a finding concordant with previous studies demonstrating increased ferritin levels in ALS patient plasma and cerebrospinal fluid (29–32). These increases in ferritin were also associated with disease progression in ALS (30, 31). Increased ferritin can be stimulated by an excess of iron and each ferritin molecule has the capability of holding up to 4500 Fe(III) ions (33). Therefore, increased ferritin could indicate increased levels of iron which have been shown to indirectly regulate TDP-43 through oxidative stress (34). In the current study, increased ferritin immunoreactivity was observed not only in cases with behavioural deficits, but also in discordant cases with no behavioural deficit but which exhibited elevated TDP-43 pathology. This suggests that ferritin accumulation may occur in the early stages of disease before behavioural symptoms occur. Susceptibility-based MRI techniques can be used to visualize and quantify iron levels, with the main source of signal on these scans coming from ferritin-bound iron (35, 36). Therefore, such MRI techniques may have potential as early biomarkers for ALS, with several studies demonstrating hyperintensities on susceptibility-weighted imaging in ALS patients that correlate with disease severity (37–39).

While iron has been shown to indirectly increase Tau phosphorylation via CDK5 and GSK3β activity, we show that there is no significant correlation between ferritin and pTau (40). Previous studies demonstrate some indirect links between iron levels and pTau, with Fe (II) appearing to mediate tau-tau interactions and structural changes (41). As iron stored in ferritin is in the Fe(III) state, it is likely that free Fe(II) would correlate more strongly with pTau levels than ferritin levels. Interestingly, one study showed that while Tau and CSF ferritin levels both independently predicted rate of cognitive decline, they did not interact in the prediction model, suggesting differential mechanisms (42).

These data clearly support the idea that protein misfolding and iron regulation therapies may hold promise in this patient population (43). Indeed, clinical trials have investigated the role of ameliorating protein misfolding in ALS patients previously, such as with arimoclomol, a heat shock protein (HSP) co-inducer that we have shown previously through meta-analysis to significantly reduce behavioural outcome measures in animal models (43). When trialed in ALS patients, arimoclomol had no effect on survival outcomes; however, this trial had no cognitive or behavioural outcome measure (44, 45). It is therefore important to note that trials of therapies specifically targeting protein misfolding that have not been successful have not included a cognitive or behavioural primary endpoint nor a pathological assessment of *ante-mortem* imaging or *post-mortem* tissue to confirm whether TDP-43 had indeed been cleared.

This underscores the importance of utilizing imaging and cognitive and behavioural endpoints that can assess non-motor brain regions, especially when evaluating the effects of therapies targeting protein misfolding.

In conclusion, our data, taken together with the developing body of literature describing extra-motor manifestations of ALS, demonstrate that specific clinical correlates of TDP-43 pathology exist in both imaging and neuropsychological profiling. These data further underscore the unmet need for personalized clinical trials using screening tools such as the ECAS, in combination with targeted neuroimaging of relevant brain regions such as the amygdala, to not only effectively stratify patients and monitor treatment, but also to potentially identify individuals in early or even pre-symptomatic states.

## Acknowledgements

The authors would like to thank (i) the MRC Edinburgh Brain Bank for supplying all *post-mortem* brain material and the Scottish MND Register/CARE-MND Consortium for all clinical and demographic data. (ii) The Scottish MND Clinical Specialist, team in discussing and obtaining consent from people with MND for inclusion in these resources. (iii) People affected by MND and their families for their hugely important and kind donation to the Brain Bank.

## Study funded by

MND Scotland (2021/MNDS/RP/8440GREG) to JMG. NIH (1R01NS127186) to JMG employing JO’S, FR and HS. Target ALS (BB-2022-C4-L2) to NS, GGT, EZ, and JMG employing FMW and MG. The Wellcome Trust (108890/Z/15/Z) to OMR.

## Author contributions

OR, FW, JO’S, FR, MG, AP, NS, GGT, EZ, HS, and JG contributed to one or all of the following: (i) conception and design of the study, (ii) acquisition and analysis of data and (iii) drafting a significant portion of the manuscript or figures.

